# Prior infection induces long-lasting partial immunity to reduce transmission within flocks in an avian host-pathogen system

**DOI:** 10.1101/2025.03.23.644804

**Authors:** Madeline C. Sudnick, Erin L. Sauer, Sarah E. DuRant

## Abstract

Population-level differences in the ability to maintain immunity can lead to contrasting survival outcomes between groups and determine how future outbreaks will spread. Pathology and infectiousness during subsequent infections can be influenced by immunity and an increase in resistance or tolerance of individuals. However, determining the longevity and effectiveness of immunity in wildlife population is challenging due to lack of infrastructure for long term monitoring of populations and individuals and host-pathogen specific tests. Further, predicting wildlife disease dynamics requires an understanding of individual-level heterogeneity in pathology and behavior as well as a knowledge of how populations with differing partial immunity will transmit disease, which is currently limited. Using an avian host-pathogen system, we ran two experiments to determine if previously exposed birds had immunity to MG after three years and if *Mycoplasma gallisepticum* (MG) transmission differed between first and second infection index birds. Birds retained partial immunity to MG for three years after their original infection and showed signs of being resistant to infection. The transmission experiment revealed that although first and second-infection index birds experienced pathology and pathogen growth, only first infection birds transmitted MG. Three years could represent lifetime protection for individuals who survive their first infection and a reduction in risk for other members of their flock. Our research has important implications for understanding MG epidemics in wild populations of birds as well as disease in wildlife more broadly.

## Introduction

Understanding the persistence and spread of disease in wildlife populations is critical, as wild populations function as reservoirs for zoonotic pathogens, and infectious diseases have posed risk to biodiversity conservation (Dulberger et al. 2010; Jolles et al. 2021; Gortázar et al. 2022; Hoyt et al. 2023). Increasing calls for elevated wildlife disease monitoring require us to understand how wildlife transmit disease within populations (Barroso et al. 2021; Cardoso et al. 2022), which involves understanding individual-level heterogeneity in pathology and behavior as well as analysis of contact networks within populations (Gilbertson et al. 2021), both of which are challenging to determine in the wild. Especially important are long-term studies, which can be challenging to conduct due to the difficulties surrounding tracking of wild living individuals over multiple years and epidemics (Barroso et al. 2021). Therefore, there is limited knowledge of how populations in various stages of an epidemic will transmit disease. We aim to help fill this gap by experimentally determining the contribution of previous exposure to disease on long-term immunity and future transmission.

Epidemic dynamics can be affected by host susceptibility and behavior, both of which can be impacted by prior infections, and can lead to immunity or partial immunity of some individuals (Cizauskas et al. 2014; Lopes et al. 2016). Many wildlife diseases can exist at low infection rates for long periods of time, then increase to epizootic levels of infection when there is an increase in susceptible individuals (Kramer-Schadt et al. 2009). Despite this, we often do not know how much naïve versus previously-exposed individuals contribute to disease transmission. Prior exposure can alter host physiology (e.g., immune function, health condition) to affect how pathogens establish infection and transmit to new hosts. Host physiology also impacts the odds of an individual surviving the infection and, more broadly, the odds of disease-driven population declines (Adelman et al. 2013b). Further, surviving infectious individuals can spread disease through social behavior while recovered individuals who retain a level of immunity post-infection may reduce transmission (Faustino et al. 2004). Thus, understanding post-infection physiological changes including disease tolerance and immunity, is critical for predicting how prior infection alters population transmission dynamics.

Transmission within populations relies on direct and indirect interactions between individuals and infection characteristics of the infected individual (Hall 2022). Social interactions such as allopreening and sharing food as well as aggressive interactions such as territorial disputes are a major factor in transmission (Sydenstricker et al. 2005; Adelman et al. 2015; Aiello et al. 2016). Individuals who have more contact time with others and have larger networks are more frequently infected (Sydenstricker et al. 2005; Aiello et al. 2016). Due to this risk, noticeably ill individuals are often avoided by conspecifics, changing the number of interactions diseased individuals have (Sydenstricker et al. 2005). Signs of infection that may lead to avoidance and transmission can be shaped by pathogen growth in hosts and host immune responses, both of which can vary with prior pathogen exposure (Cassirer et al. 2017; Rodríguez-Pastor et al. 2023) and result in resistance to or tolerance of individuals to the pathogen to affect transmission (Gates et al. 2021, Connelly 2023 (Cassirer et al. 2017; Rodríguez-Pastor et al. 2023)). Higher resistance to a pathogen leads to slower pathogen reproduction (i.e., lower pathogen loads) whereas higher tolerance emphasizes lower pathology in the individual no matter the pathogen load (Råberg 2014). Populations or individuals with high tolerance may have reduced fever, inflammation, clinical signs of illness and proinflammatory signaling while maintaining high pathogen loads (Adelman et al. 2013b). On the other hand, highly resistant individuals can have severe clinical signs and inflammation, but are in a highly infectious state (i.e., harboring high pathogen load) for a shorter period of time (Bonneaud et al. 2019; Ruden and Adelman 2021). While tolerance protects individuals from the consequences of immune activation, having high pathogen loads for longer periods of time could increase transmission likelihood as higher pathogen loads increase transmission of infection more effectively with less social interaction (Aiello et al. 2016). Tolerance can also lead to greater pathogen virulence over time through mutations as higher pathogen loads increase transmission likelihood with less social interaction (Aiello et al. 2016, 2019; Bonneaud et al. 2019; Seal et al. 2021; Ruden and Adelman 2021).

The common avian bacterial pathogen *Mycoplasma gallisepticum* (MG) is a useful study system for exploring the effects of previous infections on pathology, immunity, and transmission due to the rapid selection caused by outbreaks. MG causes severe conjunctivitis in house finches (*Haemorhous mexicanus*) and has led to population declines (Hartup et al. 2001). MG is primarily a disease of captive poultry, but in 1994 the bacteria jumped to wild house finches in the Mid-Atlantic region (Dhondt et al. 2005; Grodio et al. 2013). Following that initial epidemic, east coast house finch populations experienced drastic declines of as much as 60% (Hochachka and Dhondt 2000; Altizer et al. 2004).

MG is now established throughout much of the continental US, so the likelihood of wild birds encountering the pathogen multiple times is high. MG re-emergence is cyclic, with an autumn peak as juvenile naive birds enter the flock, and an early spring peak as winter flocks disperse and hormonal changes potentially disrupt immunocompetence (Altizer et al. 2004, Hosseini et al. 2004). Multi-year fluctuations in MG prevalence also occur in 2-3 year oscillations, leading to asynchrony in epidemics among regions (Altizer et al. 2004). Birds maintain partial resistance to reinfection through MG-specific antibodies for at least 14 months (Dhondt et al. 2017) and show lower pathology, reduced pathogen loads and faster recovery during reinfection than naïve birds (Hosseini et al. 2004; Sydenstricker et al. 2005; Dhondt et al. 2017; Leon and Hawley 2017; Fleming-Davies et al. 2018). However, we do not know yet whether those differences will translate to differences in transmission potential. Additionally, we do not currently know if the partial immunity due to resistance and tolerance lasts beyond 14 months into longer periods that match the 2-3 year MG oscillations in wild populations.

Adaptation towards resistance and tolerance over time in wild populations after the initial spread of MG complicates the relationship between bird physiology, behavior, and disease transmission (Bonneaud et al. 2019; Henschen et al. 2023). Because lesions are a result of inflammation, individuals with high tolerance often have reduced lesions, and are therefore less likely to transmit the pathogen (Ruden and Adelman 2021). However, individuals with increased resistance may have fewer copies of the pathogen to deposit and could have reduced transmission potential as well. Further, the amount of time individuals spend on feeders impacts transmission (Adelman et al. 2015), making behavior at bird feeders and interactions between conspecifics important in understanding disease dynamics. MG is spread via fomite transmission at bird feeders (Adelman et al. 2013a), and deposition of bacteria on fomites as well as transmission generally increases with eye lesion severity and pathogen load (Adelman et al. 2013a; Williams et al. 2014; Bonneaud et al. 2020), so birds with visible symptoms are more likely to transmit the pathogen.

Understanding post-infection physiological changes including disease tolerance and immunity is critical for predicting population transmission dynamics and future epidemics (Demas and Nelson 2012). Therefore, we aim to illustrate differences in disease transmission due to infection history and determine which host traits drive those differences in transmission potential. In the first experiment, we determined whether previously exposed birds still had resistance to MG after three years, which could help explain the 2-3-year fluctuations in MG epidemics in the wild (Altizer et al. 2004). We predicted that the re-infected birds would have increased resistance to MG, and therefore have lower pathogen loads and higher levels of MG-specific antibodies. We predicted that re-infected individuals would also be more tolerant of infection and have lower eye lesion severity and less severe changes in body mass and condition. In the second experiment, we assessed differences in transmission of birds experiencing their first infection against birds experiencing their second infection with MG. We predicted that birds in their first infection would have more transmission in their flocks. This work will help illustrate how pathogens can spread across populations with a mix of previously infected and naïve birds.

## Methods

### Study System

We used domestic canaries (*Serinus canaria domestica*) and MG as our host pathogen system. Canaries are a useful lab model species for MG host-pathogen dynamics because they can live long-term in the lab while exhibiting normal life history patterns and exhibit similar pathology and pathogen load when infected with MG as wild house finches (Hawley et al. 2011; Perrine et al. 2025). All experimental procedures were approved by the Institutional Animal Care and Use Committee of the University of Arkansas.

### Experiment 1 Experimental design

In the first experiment, we infected 20 birds in June through August 2020, monitored each individual until they recovered from MG, then kept them completely isolated from MG until they were re-infected in April 2023. During infection, we monitored their bacterial loads, eye lesion severity, white blood cell counts, hematocrit, body condition, and antibody presence for three weeks. Ideally, we would have included a treatment to control for age versus number of exposures, but could not due to logistical constraints. Instead, we opportunistically compiled data from other experiments where birds were first exposed to MG at ∼3 years of age to compare to 3-year-old birds experiencing a second MG infection (see supplemental material).

Following the methods of Hawley et al. (2011), we inoculated birds in their palpebral conjunctiva with 25 μL of MG inoculum diluted 5.6% in Frey’s media for a final inoculum concentration of 2.8×106 CCU/mL (VA1994; 5.0×107 CCU/mL stock). After birds were inoculated, we held them in a paper bag for 5 minutes to allow full absorption of MG into the eye.

We monitored disease pathology and pathogen load on days 0, 1, 3, 5, 7, 10, 14, 17, and 21. Disease pathology was determined by scoring conjunctival lesions on a 0–3 scale following Sydenstricker et al. (2005) where 1 = minor swelling around the eye, 2 = moderate swelling, eversion of the conjunctival tissue, and conjunctival discoloration, and 3 = the eye is nearly hidden by swelling and crusted exudates and feather loss is present around the eyes. To measure pathogen load we swabbed all birds conjunctiva for 5 seconds with a cotton swab saturated in tryptose phosphate broth (TPB). Swabs were cut and placed into a tube of 300 ml TPB, which was frozen until processing for quantification of MG load by qPCR. During this time, we also weighed birds and evaluated their condition.

On days 0, 2, 7, 14, and 21, we collected a small blood sample to use for assaying MG-specific antibodies. Blood samples were centrifuged and plasma was stored at -20°C until antibodies were quantified using ELISAs.

### Experiment 2 Experimental Design

In the second experiment, flocks of four birds (two male and two female) were divided into two treatments: all naïve to MG (n = 8 flocks) or all that were previously exposed to MG once in February 2021 (n = 9 flocks). Each flock was housed in a 77 x 46.5 x 92.5 cm cage with four perches to allow room for flight, avoidance, and social interactions. Each cage contained a single bird feeder and water bottle. Flocks were separated by sight from neighbors. We acclimated birds to flock living for a week prior to exposure (Marché et al. 2018; Haakenson et al. 2019). Before introducing birds to flocks, we banded all individuals on the leg with an external tag including a RFID reader interfaced with the bird feeder. This allowed us to determine how much time each bird was spending at a feeder. To maximize transmission likelihood, we analyzed RFID data of the last 24 hours of the acclimation period and chose the index bird by determining which male bird was at the feeder the most (Sauer et al. 2024a). After acclimation to group housing for one week, the index bird was inoculated with MG using the methods described above.

We monitored disease pathology of flock members daily by scoring conjunctival lesions. Every three days, we collected eye swabs to assess pathogen load from each individual and monitored body mass, fat, and hematocrit. The eye score and qPCR measurements were used to determine when and if transmission occurred in flocks. Initial transmission was defined as the first day on which any non-index bird had an eye score or pathogen load above 0 (Adelman et al. 2015). Every 6 days we collected a blood sample for quantifying MG-specific antibodies. We also collected the number of seconds each bird was on the feeder each day using the RFID tags and receiver. We monitored all for 28 days, after which the birds were placed into individual cages until 60 days post-infection for birds with MG or 60 days post infecting the index bird for individuals without MG.

### Quantifying pathogen load and MG antibodies

In both experiments, we used a QIAGEN DNeasy Blood and Tissue kit (Qiagen, Valencia, CA) to extract bacterial DNA from eye swab samples taken from all birds. We then determined pathogen load using quantitative PCR following the procedure outlined by (Grodio et al. 2008) that targets the mgc2 gene of MG. Final concentrations were calculated by multiplying 3 μl, the amount of DNA sample used, by 66.666, to be comparable to the 200 μl that was produced from the elution step.

In both experiments, MG-specific serum antibodies were quantified using the IDEXX MG antibody enzyme-linked immunosorbent assay test kit (IDEXX, Cat#99-06729; Grodio et al. 2009). A blocking step was added to the original assay kit’s protocol, with the addition of 300 μL of 1% bovine serum albumin (Pierce 10X BSA; Thermo Fisher Scientific) in phosphate-buffered saline to room temperature plates before they were incubated. All plates were washed three times with phosphate-buffered saline containing 0.05% Tween 20. Serum samples were diluted 1:50 in sample buffer and were then plated to be run in duplicate. Intensity of light absorbed by the serum samples was measured at 630 nm using a spectrophotometer and an ELISA value was then calculated.

### Statistical analysis

All analyses were conducted in RStudio using R V 4.4.3 (R Core Team 2025).

Experiment 1: We used generalized linear models and generalized additive models to determine differences in eye lesion scores, pathogen load, antibody levels, and tolerance (Wood 2023). We log transformed pathogen loads to meet analysis assumptions. To evaluate changes in body condition due to each infection, we calculated change in mass, hematocrit and fat during the experiment. We used repeated measures ANOVAS to determine differences in the infections while controlling for the non-independence from the same individuals going through both the first and second infections. To determine if differences in first and second infections were the result of the bird’s increased age, we conducted a supplementary analysis comparing eye scores, pathogen loads, MG-specific antibodies, and hematocrit levels in secondary exposures of the individuals in this study to those in first exposures of similar aged individuals (3 yo) in other studies (see supplementary methods for details).

Experiment 2: To determine the difference in transmission between the previously MG-infected and MG-naive groups, we used the Fisher’s exact test. To determine the impact of host traits and disease severity on whether an index bird transmitted MG, we conducted a generalized mixed model with a binomial distribution and used model selection to determine which predictors to include in the final model (Barton 2019; Pinheiro et al. 2021). We included only first infection index birds in these models because no second infection index birds transmitted MG. We calculated area under the curve for variables that changed over time and only included the first 10 days of infection, as this is when the birds were transmitting to other members of their flock. We also calculated maximum value for antibody presence to consider the highest level of learned immune response the index bird had, as antibody presence does not return to base levels within the time of the experiment. Predictor variables in the global model included bacterial loads (AUC), eye lesion severity (AUC), seconds at the feeder (AUC), hematocrit (AUC), mass (AUC) and antibody level (max). We transformed variables with non-normal distributions to meet the assumptions of the model.

To determine whether there were physiological variables that predicted whether a non-index bird would become infected we conducted a generalized linear model with a binomial distribution and used model selection to determine which physiological variables of non-index birds to include in the final model. We calculated area under the curve for variables that changed over time, and day zero value for variables that it was most important to have the pre-exposure value to only include the time where the individual could become infected. Predictor variables in the global model included time spent at the feeder (AUC), condition (fat, mass, hematocrit) and antibody amount. We included the random effect of bird ID to avoid pseudoreplication.

## RESULTS

### Experiment 1

All 20 birds in our study presented with lesions in both their first and second infections, with peak lesion severity occurring in days 5-7. All birds returned to a lesion severity of zero by the end of our study on day 28. Birds in their first infection recovered by day 20, whereas birds in their second infection recovered from their eye lesions by day 15. We found that second infection birds had lower eye lesion severity (t = -1.95, *p* = 0.05; Figure 1A) and pathogen loads (χ² = 5.35, *p* = 0.02; Figure 1B). Antibody presence increased over the course of the infection in both first and second infections, leading to higher antibody values overall during the second infection (χ² = 49.36, *p* < 0.001; Figure 1C). Birds had slightly higher tolerance during the second infection, however, the difference was not significant (t = 1.41, *p* = 0.16; Figure 1D).

**Figure 1.**
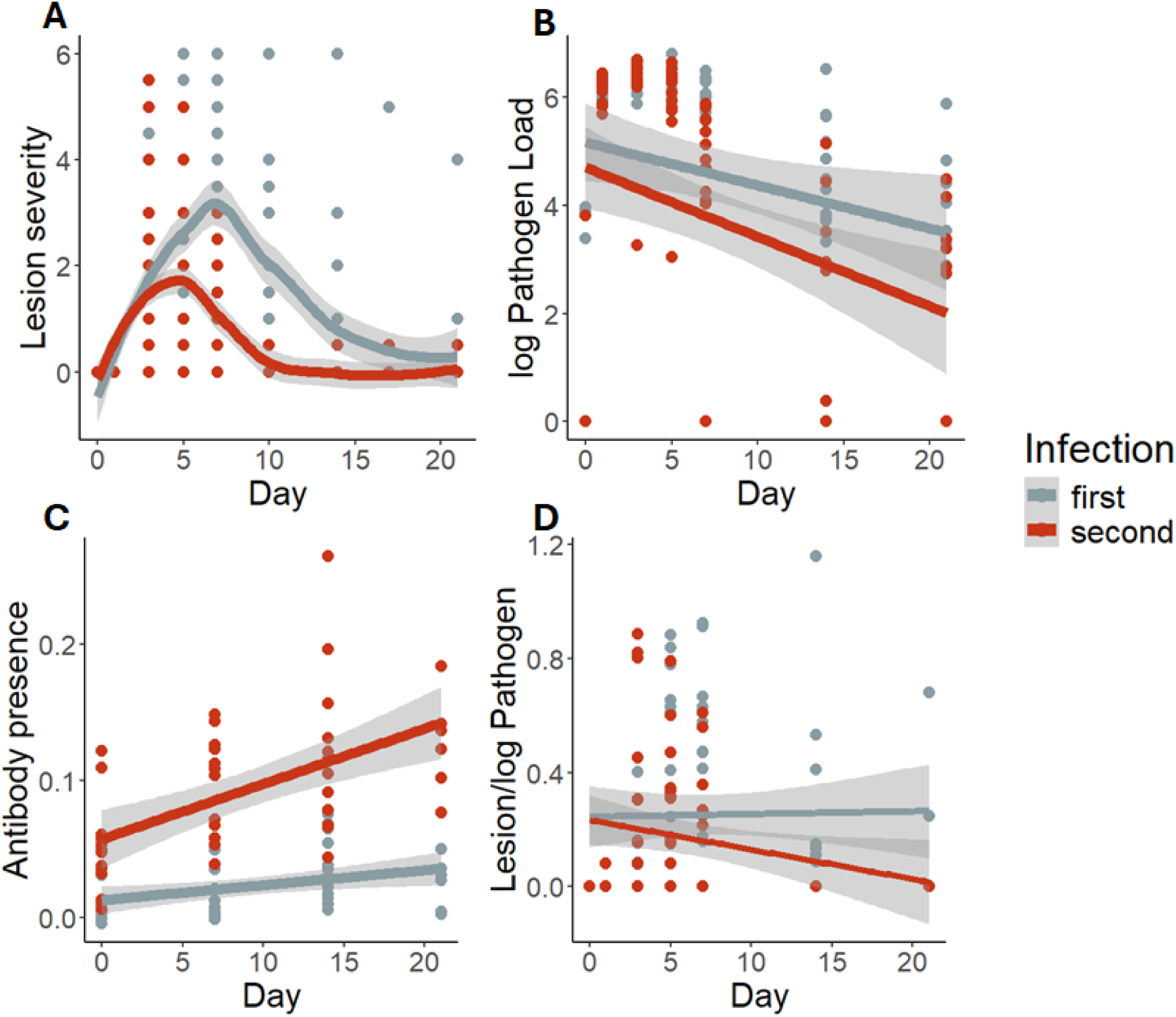
Domestic canaries were re-infected with Mycoplasma gallisepticum three years after an initial infection. A) Conjunctival lesion severity was monitored for 28 days, with each eye ranked on a 0-3 severity scale, totaling to a possible maximum of 6. Birds in their second infection had lower eye lesion severity than birds in their first infection (*t* = -1.95, *p* = 0.05). B) Pathogen load was monitored for 28 days by using qpcr to count the copies of bacterial DNA taken from an eye swab. Birds in their second infection had lower pathogen loads than birds in their first infection (χ² = 5.35, *p* = 0.02). C) Antibody presence was monitored for 28 days by using immune assays to count the copies of MG-specific IgY antibodies present in blood plasma. Antibody presence was maintained during the three year period without infection, and was raised before inoculation of the second infection. Birds in their second infection had higher antibody presence than birds in their first infection (χ² = 49.36, *p* < 0.001). D) Tolerance was monitored for 28 days by dividing lesion severity by the pathogen load. This demonstrates how birds in each infection respond differently in their inflammation to similar pathogen loads. Birds in their second infection had lower lesion severity with similar pathogen loads, however, there was no significant difference in tolerance (t = 1.41, *p* = 0.16).

Birds had greater mass loss during their second infection (*F* = 6.78, df = 19, ges = 0.13, *p* = 0.02; Figure 2). There was no difference in change in hematocrit (*F* = 0.75, df = 19, ges = 0.02, *p* = 0.40), or change in fat score (*F* = 1, df = 19, ges = 0.02, *p* = 0.3) between first or second infections.

**Figure 2.**
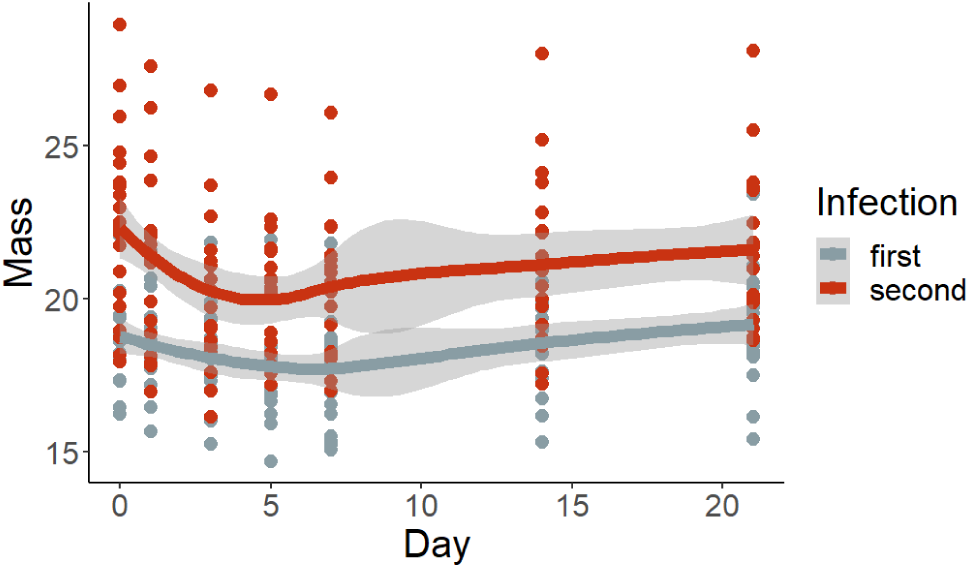
Domestic canaries were re-infected with *Mycoplasma gallisepticum* three years after an initial infection. Mass was monitored for 28 days. Mass decreased during peak infection but returned to similar values by the end of infection. Birds in their second infection lost more mass during peak infection than birds in their first infection (F = 8.493, df = 12, ges = 0.259, *p* = 0.013).

### Experiment 2

We found that only first-infection birds transmitted MG to any other member of their flock (*p =* 0.009; Figure 3). Five of the eight first-infection flocks had transmission to at least one non-index bird. Four flocks had multiple transmission events, and one had a single transmission event. Of the 15 non-index birds within flocks that had at least one transmission event, 12 presented with either eye lesions or copies of MG bacterial DNA.

**Figure 3.**
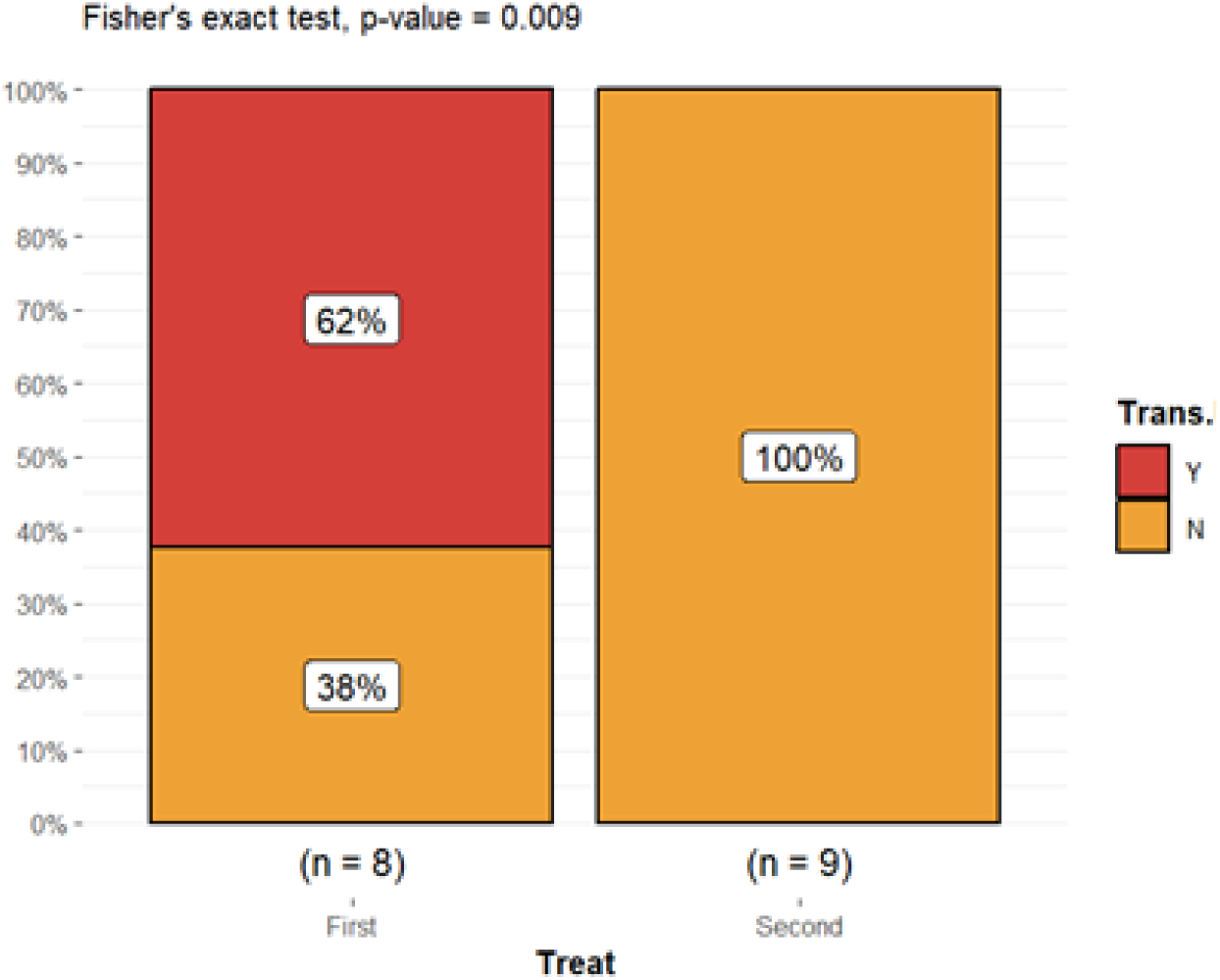
Bar graph showing Fisher’s exact test comparing transmission in canaries (*Serinus canaria domesticus*) experiencing their first or second infection with *Mycoplamsa gallisepticum* (MG) (p = 0.009). Flocks of either four naïve birds (n=8) or four previously MG-infected birds (n= 9) were housed together for 7 days. After 7 days one bird from each flock was infected with MG (index bird) and flock mates were monitored for signs of infection, i.e., that transmission had occurred, for 28 days.

We found that first infection birds had higher pathogen load (β = -2.31 ± 0.55, df = 15, *t* = -4.20, *p* < 0.001), higher eye lesion severity (β = -1.31 ± 0.49, df = 15, *t* = -2.67, *p* = 0.02), and lower antibody presence (β = 0.026, ± 0.01, df = 15, *t* = 3.32, *p* = 0.005) than second infection flocks. Birds did not differ between treatments for tolerance (β = -0.12 ± 0.08, df = 15, *t* = -1.55, *p* = 0.14), hematocrit (β = -4.96 ± 2.73, df = 15, *t* = -1.81, *p* = 0.089), mass (β = 1.57 ± 1.12, df = 15, *t* = 1.40, *p* = 0.18) or number of seconds spent on the bird feeder (β = -0.42 ± 0.33, df = 15, *t* = -1.25, *p* = 0.23).

When exploring what traits of index birds predicted transmission, we found that our top models included the null model and the model including only hematocrit (ΔAICc = 0.3; Table 3). In that model, hematocrit was not significantly different between birds that did and did not transmit infection (β = 0.28 ± 0.18, *t* = 1.55, *p* = 0.12; Figure 4; Table 2).

**Figure 4.**
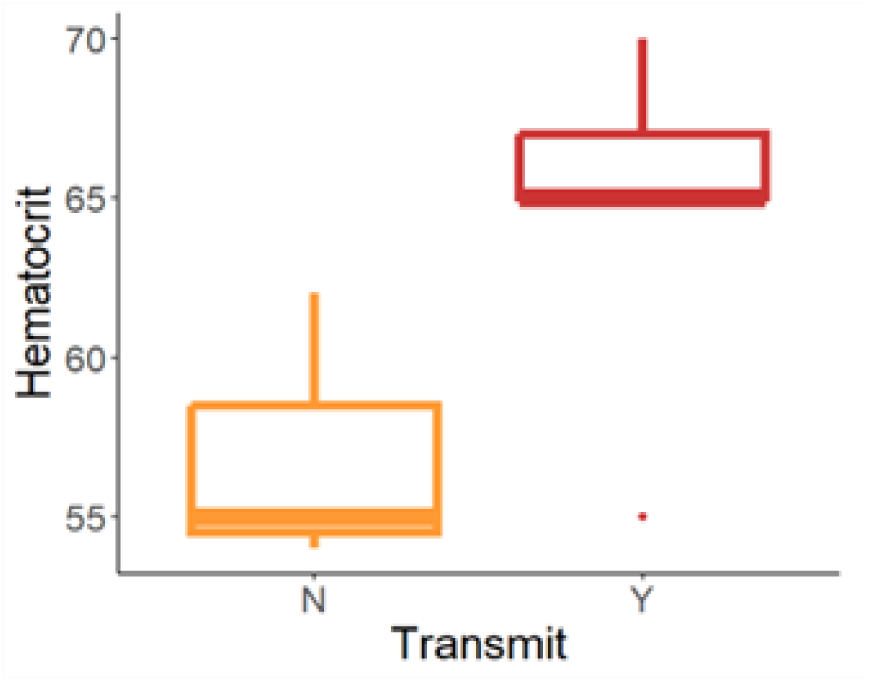
Box plot comparing hematocrit in canaries (*Serinus canaria domesticus*) that either did (n = 5) or did not (n = 3) transmit *Mycoplamsa gallisepticum* (MG) during their first infection. To measure hematocrit, we measured the packed red blood cell volume in a blood sample from each bird (β = 0.278 ± 0.1799, t = 1.546, *p* = 0.122). Flocks of either four naïve birds (n=8) or four previously MG-infected birds (n= 9) were housed together for 7 days. After 7 days one bird from each flock was infected with MG (index bird) and flock mates were monitored for signs of infection, i.e., that transmission had occurred, for 28 days.

**Table 1.**
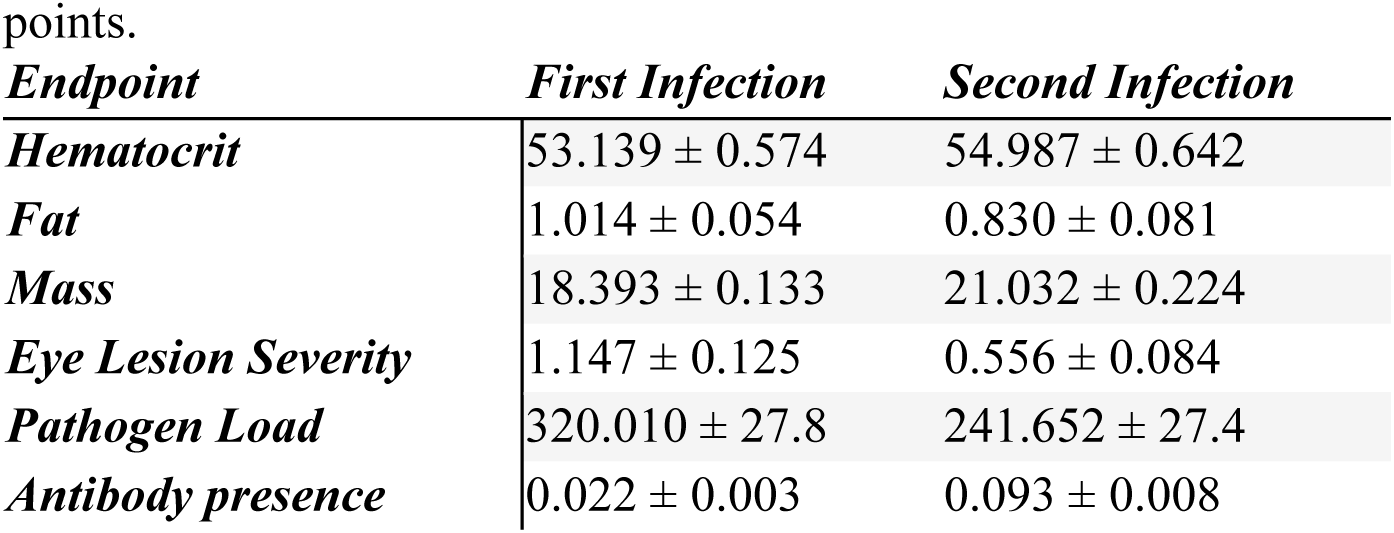
The mean ± standard error of condition and pathology endpoints measured in domestic canaries that were re-infected with *Mycoplasma gallisepticum* three years after an initial infection. We monitored condition and pathology for 28 days and averaged endpoints across all sampling time points.

**Table 2.** Model selection table comparing transmitting and non-transmitting first infection index birds (n = 8). The models compared included all combinations of the predictor variables in the global model including bacterial loads (AUC), eye lesion severity (AUC), seconds at the feeder (AUC), hematocrit (AUC), Tolerance (AUC), mass (AUC), and antibody presence (max). Top models were considered those with Δ AICc under 2. Models presented in table are Δ AICc under 5. The top models included only hematocrit and the null model.

| ***Predictors*** | ***AICc*** | ***$\Delta$ AICc*** |
| --- | --- | --- |
| ***Null*** | 13.3 | 0 |
| ***Hematocrit*** | 13.6 | 0.33 |
| ***Pathogen Load AUC*** | 15.6 | 2.38 |
| ***Feeder Visits AUC*** | 15.6 | 2.38 |
| ***Mass AUC*** | 16.5 | 3.21 |
| ***Tolerance AUC*** | 16.8 | 3.59 |
| ***Antibody presence max*** | 16.9 | 3.66 |
| ***Eye Lesion Severity*** | 17 | 3.73 |

**Table 3.** Model selection table comparing receiving and non-receiving non-index birds in flocks with at least one transmission event (n = 15). The models compared included all combinations of the predictor variables in the global model including time spent at the feeder (AUC), fat, hematocrit, mass, and antibody presence. Top models were considered those with Δ AICc under 2. Models presented in table are Δ AICc under 5. The top model included only mass.

| *Predictors* | *AICc* | *$\Delta AICc$* |
| --- | --- | --- |
| *Mass* | 12.9 | 0 |
| *Mass + Antibody Presence* | 16.7 | 3.79 |
| *Mass + Hematocrit* | 16.9 | 4 |
| *Mass + Feeder Visits AUC* | 17.1 | 4.17 |

Our analyses of traits of non-index birds that were in flocks with at least one transmission event (n = 15) revealed that the top model included only mass of the bird (ΔAICc other models >2. Mass was lower in birds that received the infection than those that did not (β = -18.32 ± 2.06, df = 13, *t* = -8.90, *p* < 0.001; Figure 5, Table 3).

**Figure 5.**
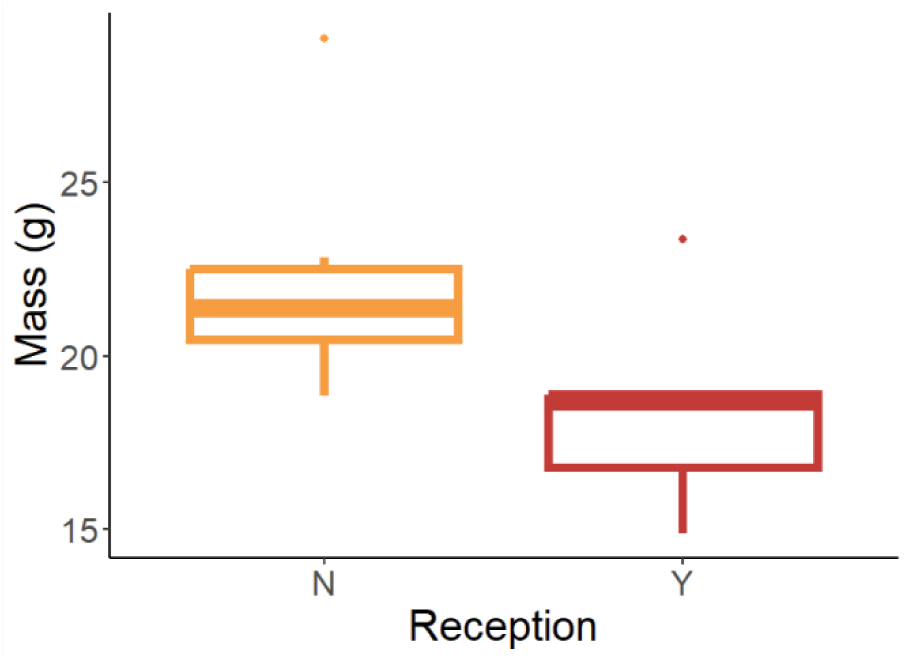
Box plot comparing mass in non-index canaries (*Serinus canaria domesticus*) that either did or did not receive *Mycoplamsa gallisepticum* (MG) during the experiment. Receiving birds (Y) weighed less than non-receiving birds (N) in flocks where transmission occurred (β = - 18.318 ± 2.057, df = 13, *t* = -8,904, *p* < 0.001). Flocks of either four naïve birds (n=8) or four previously MG-infected birds (n= 9) were housed together for 7 days. After 7 days one bird from each flock was infected with MG (index bird) and flock mates were monitored for signs of infection, i.e., that transmission had occurred, for 28 days. We measured the mass of the non-index birds on the day of infection.

## Discussion

We investigated the effects of previous infection on long-term retention of immunity, development of resistance to MG, and disease transmission. We found that canaries retained partial immunity to MG for three years after first infection and that birds develop increased resistance to MG infection. As many songbirds, including house finches, may only live for a few years after their infection in the wild due to general causes of mortality (Veit and Lewis 1996, Adelman et al. 2017), three years could represent lifetime protection for individuals who survive their first infection, and therefore lifetime reduction in disease transmission within their flock.

Our transmission experiment revealed that although both first and second-infection index birds became infected and experienced pathology, only first infection birds transmitted infection indicating that first infection birds will contribute more to disease spread and infection dynamics. Long term immunity to MG may also reduce indirect mortality of MG in birds experiencing second infections. Reinfected birds had less severe lesions, which should lower their predation risk because severe lesions obfuscate approaching threats to reduce anti-predator behavior (Adelman et al. 2017). Therefore, total population loss to MG both directly and indirectly is likely reduced during repeat infections.

We found evidence that canaries had increased resistance to MG during second infections. Second infections led to lower total pathogen loads, driven by the more rapid clearing of infection post peak, showing increased resistance to MG. Age could contribute to the lower pathology and pathogen growth in second infection birds, but older birds experiencing their first infection had more severe disease outcomes than second infection birds of the same age, indicating that naivete and not age are diving the differences in disease outcomes we detected in the first experiment.

Our findings of increased resistance in previously MG-exposed canaries are consistent with studies of house finches in wild populations exposed to MG, which have been found to increase in both resistance and tolerance with time post introduction of the pathogen (Adelman et al. 2013, Bonneaud et al. 2019, Henschen et al. 2023). For instance, a study comparing house finch populations showed faster clearance of bacteria during second infections indicating resistance to the pathogen (Bonneaud et al. 2019). They have also been found to rapidly evolve tolerance within 25 years, with MG exposed populations having lower eye lesion severity even at similar pathogen loads (Henschen et al. 2023). Taken together, our results along with these other studies indicate that populations have primarily been found to be unified in their tendency towards resistance and tolerance, likely due to the strains of MG they encounter.

Resistance in our study may have improved through heightened MG-specific antibody presence. On the day of inoculation MG specific antibodies were lower in birds experiencing their first infection than those experiencing their second infection, suggesting that after an initial infection, canaries maintain the production of antibodies for at least three years. Similarly, MG-specific antibody response to an inoculation is higher in house finch populations that have been previously exposed to MG (Gates et al. 2021). The maintenance of MG-specific antibodies could also contribute to tolerance as could activation of fewer immune genes. A previous study found that activation of immune genes prior to exposure led to increased tolerance in canaries, and these birds activated fewer genes during infection than less tolerant birds (Sauer et al 2024b).

Populations of house finches with increased tolerance had more targeted immune responses early in infection and a reduction in up-regulated pro-inflammatory genes early in infection (Henschen et al. 2023). The decrease in pro-inflammatory response could be linked to birds having lower eye inflammation during their second infection Long-term changes in host responses to MG could account for the differences in transmission we found between first and second infection birds. Like the findings in experiment one, we found differences in disease outcomes between first and second infection birds in the transmission experiment, with first infection birds having higher pathology, higher eye lesion severity and lower specific antibody presence. These findings indicate that second infection individuals maintained partial immunity during the time between their first and second inoculation. Although we saw a complete elimination of transmission in second infection birds, these results differ from a previous study that found that asymptomatic second infection adults can transmit MG in flocks (Dhondt et al. 2012). They found that while the presence of naïve juveniles or the introduction of MG is more important to an epidemic than time of year, asymptomatic recovered adults can initiate the epidemic. The differences in our results could potentially be due to species difference or host age. While canaries are a good captive model for MG due to their similar presentation of infection in the conjunctival lesions, there may be slight differences from house finches in their immunity and how they respond to reinfection. Dhondt et al. (2012) also explored reinfection dynamics in terms of age and seasonal changes, as the introduction of naïve juveniles can lead to disease spread, whereas we did not separate birds by age. Our results suggest that the more important aspect is naivete rather than age, and naïve birds that cross populations may lead to increased disease transmission. As MG spreads to additional avian species, as is being reported, it will be critical to observe any differences in reinfection dynamics to predict potential population declines caused by MG outbreaks (Dhondt et al. 2014).

Differences in transmission in our system could result from differences in tolerance between naïve and second infection index birds. The presence of visible eye lesions varied among index birds that were positive for pathogen load, and pathogen load was not uniformly low in birds without eye lesions. We found that eye lesion severity was higher in index birds that transmitted, however, that result was driven by lower severity eye lesions in second infection birds and was not significant in our first infection index bird model selection. Although previous studies have shown a close link between eye lesion score and pathogen load within population (Adelman et al. 2015), evidence indicates that some minimum degree of pathology is needed for transmission to occur (Bonneaud et al. 2020; Ruden and Adelman 2021). This would be consistent with what we found, as first infection birds experienced greater pathology and were the only index birds to transmit MG to flock mates. Different populations can differ in tolerance to MG, with some groups having more severe eye lesion presence at similar pathogen loads (Adelman et al. 2013b; Bonneaud et al. 2019), but it is unclear whether these differences could be attributable to variation in MG-naiveté across populations. We did not find a difference in tolerance between transmitting and non-transmitting birds, so eye lesion severity alone may be more important for transmission than the amount of bacteria in the eye at a given eye lesion severity. This may contribute to why second infection birds did not transmit to any members of their flock in our study.

When we explored impacts of heterogeneity in behavior or physiology on transmission, we found that only the null model and the model with hematocrit were included in the top models, however, there was no significant difference in hematocrit between transmitting and non-transmitting birds. The relationship between hematocrit levels and disease outcomes is variable, and likely disease specific, and can be unassociated (Granthon and Williams 2017; Names et al. 2022), negatively associated (Dawson and Bortolotti 1997) or positively associated (Booth and Elliott 2002) with disease outcomes. Papers linking hematocrit to disease typically measure hematocrit after individuals have been inoculated with a pathogen, and therefore there is a gap in understanding of how pre-infection hematocrit markers could alter transmission. The ability to fight off infection could potentially be increased by higher oxygen carrying capacity. The null model being the top model may show that there are other aspects of behavior or physiology we did not include in monitoring that contribute to transmission.

We also investigated the impact of the physiology and behavior of uninfected flock mates on the likelihood of becoming infected and found that the top model included only body mass. We compared birds that did and did not receive infection within flocks with at least one transmission and found that mass was lower in birds that received MG. Health attributes such as body condition could influence disease infection, progression, and transmission, for instance, wildlife may rely on fat stores for energy to fight off infection (Cheng et al. 2019; Perrine et al. 2025). However, individuals in good health may have improved host competency than those in reduced condition and therefore be more likely to shed pathogens (Arsnoe et al. 2011). Mass also has been linked to T-cell mediated immune response (Alonso-Alvarez and Tella 2001), so individuals with greater mass may have better immune systems more capable of fighting off infection.

While not in our top models, time at a communal feeding source was implicated as important to transmission in both of our models and previous work has linked transmission to the amount of time birds spend on the birdfeeder. Feeder visits are linked to food needs and social behaviors in our captive canaries. MG infected birds show increased sociality across variation in disease severity or pathogen loads (Langager et al. 2023), meaning that the birds with highest shedding rate could have been spending more social time on the feeder, leading to high amounts of bacteria on fomites. This is supported by studies showing that feeder use is tied to likelihood of transmission (Adelman et al. 2015; Moyers et al. 2018). Differences in disease reception and shedding among species may have led to differences in the results of our study compared to prior studies. We also may not have detected a strong significant effect because of small sample size, and future research with larger flock housing capacity could investigate differences in these behaviors. Nonetheless, anthropogenic feeding should continue to be an important focus of wildlife disease spread (Thompson et al. 2008; Sorensen et al. 2014; Murray et al. 2016), including management recommendations that increase awareness around community feeding disease spread, fomite transmission, and the importance of disinfecting surfaces wildlife come into contact with on a regular schedule (Boyd et al. 2014; Feliciano et al. 2018).

Wildlife diseases have been an important conservation and human health issue in recent years, especially with the increase of zoonotic disease outbreaks (Ferreira et al. 2021). As we see increased numbers of disease outbreaks in wild populations, it is important to understand which individuals are the most likely to transmit infection. If we can recognize at risk areas, we can predict population changes and introduce management decisions that could reduce transmission (Almberg et al. 2022). For MG, previous work has shown that lesion severity is linked to likelihood of transmission, so our results showing a decrease in inflammation during second infections suggest that transmission would be reduced in exposed populations for at least three years after an initial epidemic. Thus, epidemic scale outbreaks are unlikely during this period, until critical numbers of naive individuals enter the population via births and/or immigration and replace immune individuals that have died of emigrated (Dhondt et al. 2012). This finding mimics patterns of MG re-emergence in wild populations (Altizer et al. 2004; Hosseini et al. 2004). Previous large scale flock studies showed that previously infected birds are still vulnerable to re-infection from new outbreaks brought in by naïve individuals (Dhondt et al. 2012). However, combined with our findings, these reinfected birds are unlikely to contribute significantly to further MG transmission. Based on these findings, we would expect to see the most transmission and the most severe disease in newly established areas, such as in western states and Hawaii (Henschen et al. 2023). Understanding how long-term immunity will influence population survival is important, especially when deciding how to implement disease prevention measures. Therefore, future research should include population disease modeling where the gap between outbreaks is varied, and the responses predict which populations are at risk based on recent outbreaks.

## Acknowledgements

We thank W. Perrine and W. Kirkpatrick for their support in the lab. A. Love collected data used in analyses. J. Adelman provided advice on experimental design, statistics and RFID tags.

## Notes

https://github.com/mcsudnick/Sudnick.et.al.Mg.Infection.git

https://github.com/erinsauer/Perrine-et-al-MG-diet

### Competing Interest Statement

The authors have declared no competing interest.

