## Supplemental Methods for "Prior infection induces long-lasting partial immunity to reduce transmission within flocks in an avian host-pathogen system"

Supplementary methods

We made opportunistic use of data in the lab to determine whether age or re-infection drove the changes in disease outcomes we detected in the first experiment. First infection canaries used in this analysis were compiled from six experiments, including 3 birds used in experiment 2 (transmission experiment) of the current manuscript. Disease endpoints in these birds were then compared with disease endpoints of 3-year-old birds in experiment one when experiencing their second infection. We compiled data from experiments that were conducted between 2020 and 2024 and only one of the data sets is currently published (Weston et al. 2025). Birds were exposed to the same pathogen strain and had similar sampling regimes as used in the current experiment. Raw data are provided at {https://github.com/mcsudnick/Sudnick.et.al.Mg.Infection.git }. We used generalized linear models and generalized additive models to determine differences in eye lesion scores, pathogen load, antibody levels, tolerance, and changes in mass, fat and hematocrit over the course of the infection. We log transformed pathogen loads to meet analysis assumptions

In comparing 3year old birds in their first and second infections, we found that birds in their second infection had lower eye scores ($t$ = -6.01, $p < 0.001$), higher tolerance ($t$ = -5.896, $p < 0.001$), lower pathogen loads ( $\chi^2 = 86.01$, $p < 0.001$), higher antibody levels ( $\chi^2 = 33.89$, $p < 0.001$), and lost less fat ( $\chi^2 = 4.80$, $p = 0.03$) and less mass during infection ( $\chi^2 = 9.62$, $p = 0.001$). They also had lower hematocrit during their second infection ($\chi^2 = 4.60$, $p = 0.03$).
